# NAPR: a cloud-based framework for neuroanatomical age prediction

**DOI:** 10.1101/099309

**Authors:** Heath R. Pardoe, Ruben Kuzniecky

## Abstract

This paper describes NAPR, a cloud-based framework for accessing age prediction models created using machine learning-based analysis of neuroimaging data. The NAPR service is provided at https://www.cloudneuro.org. The NAPR system allows external users to predict the age of individual subjects using their own MRI data. As a demonstration of the NAPR approach, age prediction models were trained using healthy control data from the ABIDE, CoRR, DLBS and NKI Rockland neuroimaging datasets (total N = 2367). MRI scans were processed using Freesurfer v5.3. Age prediction models were trained using relevance vector machines and Gaussian processes machine learning techniques. NAPR will allow for rigorous and transparent out-of-sample assessment of age prediction model performance, and may therefore assist in the translation of neuroimaging-based modelling techniques to the clinic.

## Introduction

A number of recent studies have applied machine learning techniques to human functional or structural neuroimaging data to predict an individual’s age [1–6]. Differences between predicted and chronological age have been associated with a number of neurological disorders or conditions, suggesting that age estimation using MRI may have potential utility as a marker for health outcomes [4, 7–10]. In order for neuroimaging-based age prediction to be clinically useful in individual subjects, predictive models need to be accurate. A simple and effective method for robust assessment of model accuracy is out-of-sample evaluation of model performance by external users.

In this study we present a cloud-based framework for neuroimaging-based age prediction, named “NAPR”: Neuroanatomical Age Prediction using R. The NAPR system provides an interface for external users to apply predictive models to their own neuroimaging data. Users upload morphometric estimates to a cloud-based Amazon Web Services (AWS) instance running the statistical software package R via OpenCPU server [11, 12]. Morphometric estimates are input to an age prediction model which estimates the age of the subject and returns the age estimate to the end user. We believe that this system for model distribution will facilitate improvements in the accuracy of MRI-based age prediction techniques by allowing for transparent and robust out-of-sample model evaluation. The NAPR system is also readily amenable to incorporating new predictive models. An additional benefit is that external groups lacking resources to develop and implement their own models in-house can apply neuroimaging-based age prediction to their own datasets.

The NAPR web service has been provided at https://www.cloudneuro.org. We have provided a simple client script that can be used by external users to obtain age predictions on their own Freesurfer-processed data. The client script, training data and code for building and evaluating the age prediction models, and the R package for returning age predictions are provided at https://github.com/hpardoe/napr. As an example of the NAPR system, we have provided age prediction models that have been trained using Freesurfer-based cortical thickness estimates obtained from healthy control scans from the Autism Brain Imaging Data Exchange (the original ABIDE study and ABIDE II, [13]), Consortium for Reproducibility (CoRR, [14]), Dallas Lifespan Brain Study (DLBS, [15]), and NKI Rockland ([16]) datasets. We compare the performance of two machine learning regression techniques for predicting age, (i) relevance vector machines [17] and (ii) Gaussian processes [18]. We also investigated the use of undersampling as a technique for dealing for class imbalance in our training datasets; in our case, there were considerably more younger participants (age < 40 years) available than older subjects. Model performance was evaluated by measuring (i) mean absolute error for each model, and (ii) investigating the distribution of residual errors as a function of age.

## Methods

### Description of imaging datasets used in the study

Whole brain T1 weighted MRI scans of healthy controls from the ABIDE, ABIDE II, CoRR, DLBS, and NKI Rockland datasets were used in our study. Image acquisition parameters and additional details about these datasets can be found at http://fcon_1000.projects.nitrc.org. Age prediction models were trained using (i) all available healthy control imaging data, restricted to using a single scan per subject in the case of studies where multiple scans were obtained per subject, and (ii) a reduced sample, in which subjects were randomly sampled in 5-year age intervals in groups with age ranges that were overrepresented in the complete pooled dataset (ie. participants aged < 40 years).

### Image processing

Structural MRI scans were processed using the Freesurfer v5.3 default processing stream [19]. Cortical thickness surface maps were coregistered to the “fsaverage4” template, following a similar approach to that presented in [1]. The fsaverage4 template has 2562 vertices per hemisphere, yielding a total of 5124 features that were used to train the age prediction model. 200 participants were randomly selected and held out as a test dataset to evaluate model performance.

### Age prediction model training

Age prediction models were built using (i) the relevance vector machine regression method [17], and (ii) Gaussian processes regression [18], as implemented in the *kernlab* R package [20]. Models were trained on the full training sample and the undersampled dataset (referred to as “reduced” in remainder of the text) for each machine learning method, yielding four models in total: (1) *rvm.full*, (2) *rvm.reduced*, (3) *gausspr.full*, and (4) *gausspr.reduced.* The models were trained using a radial basis kernel function (“rbfdot”) and kernel hyperparameters were calculated automatically (kpar = “automatic”). Model performance was assessed by calculating the mean absolute error (MAE) of age predictions in the test dataset. Residuals (difference between predicted and chronological age) as a function of chronological age were examined to determine how well the models performed across age ranges.

### NAPR: cloud-based age predictive modelling

Opencpu version 1.6 [11] was installed on an AWS EC2 instance. An R package containing the age prediction model and a function for returning age predictions was developed on a local desktop machine and uploaded and installed on the opencpu server. External users interact with the opencpu server using a bash script that (i) concatenates left and right hemisphere cortical thickness surfaces into a tar archive, (ii) securely uploads the file to the opencpu server using the curl software package, (iii) inputs the surfaces to the age prediction model, and (iv) returns the predicted age of the participant. Performance of the NAPR system was evaluated by measuring the time taken to return age predictions for Freesurfer-based morphometric estimates from one hundred random subjects stored on a local desktop machine at the NYU Langone Medical Center and sequentially input to NAPR using the provided client program.

## Results

The full MRI dataset consisted of 2367 healthy controls obtained from the ABIDE I and II, CoRR, DLBS and NKI Rockland datasets. 200 participants were randomly selected and held out as a testing dataset for evaluating the age prediction model performance, leaving 2167 subjects as the training dataset for the complete sample. Undersampling younger subjects yielded a dataset consisting of 882 subjects (Figure 1A). The distribution of residual errors (difference between predicted and chronological age) for each machine learning approach is shown in Figure 1. The relevance vector machine models had lower mean average error than both gaussian processes models, with small improvements in MAE when full datasets were used compared with the reduced datasets (Figure 2). Although undersampling led to a higher overall MAE for both the relevance vector machine models and the Gaussian processes models, we observed lower residual errors for subjects aged greater than 40 years for both undersampled modelling approaches (*rvm.reduced* and *gausspr.reduced* models). A consistent variable distribution of residuals was observed across all four models, with younger subjects having a systematically overestimated age, and older subjects had a systematically underestimated age. Average processing time for age prediction using the cloud-based NAPR system was 3 min 14 seconds for 100 cases, or 1.94 seconds per subject.

**Figure 1:**
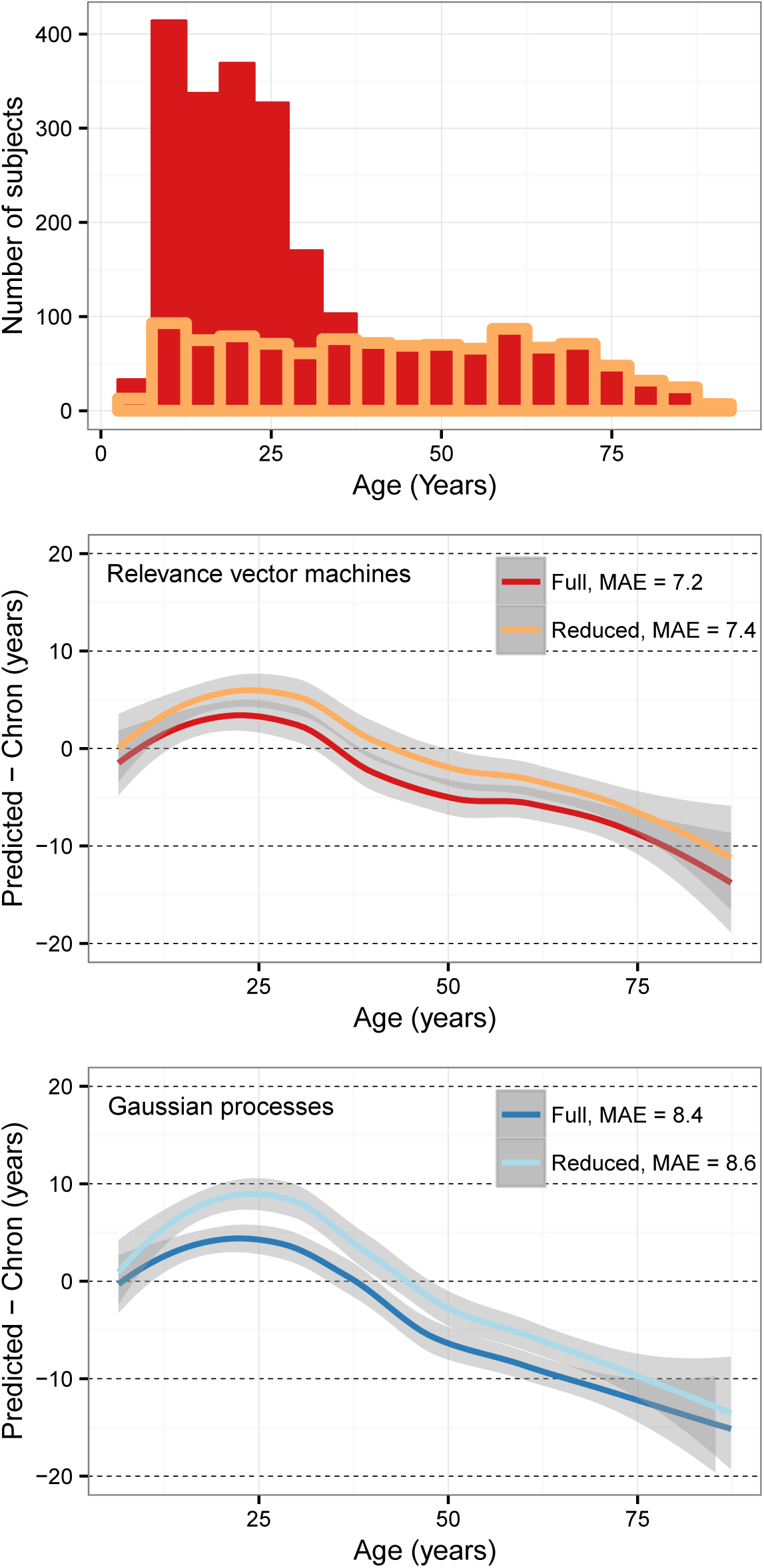
The top histogram shows the age distribution in the full sample (N = 2367) and the reduced sample (N = 1082), in which younger participants are undersampled to produce a uniform age distribution. Residual differences between predicted and chronological age show a consistent pattern independent of the machine learning method used, with systematically overestimated age in younger subjects (age < 40 years) and underestimated age in older subjects. Although overall MAE is increased when using the undersampled dataset, there appears to be a modest improvement in residuals in older subjects at the expense of higher residuals in the younger participants.

**Figure 2:**
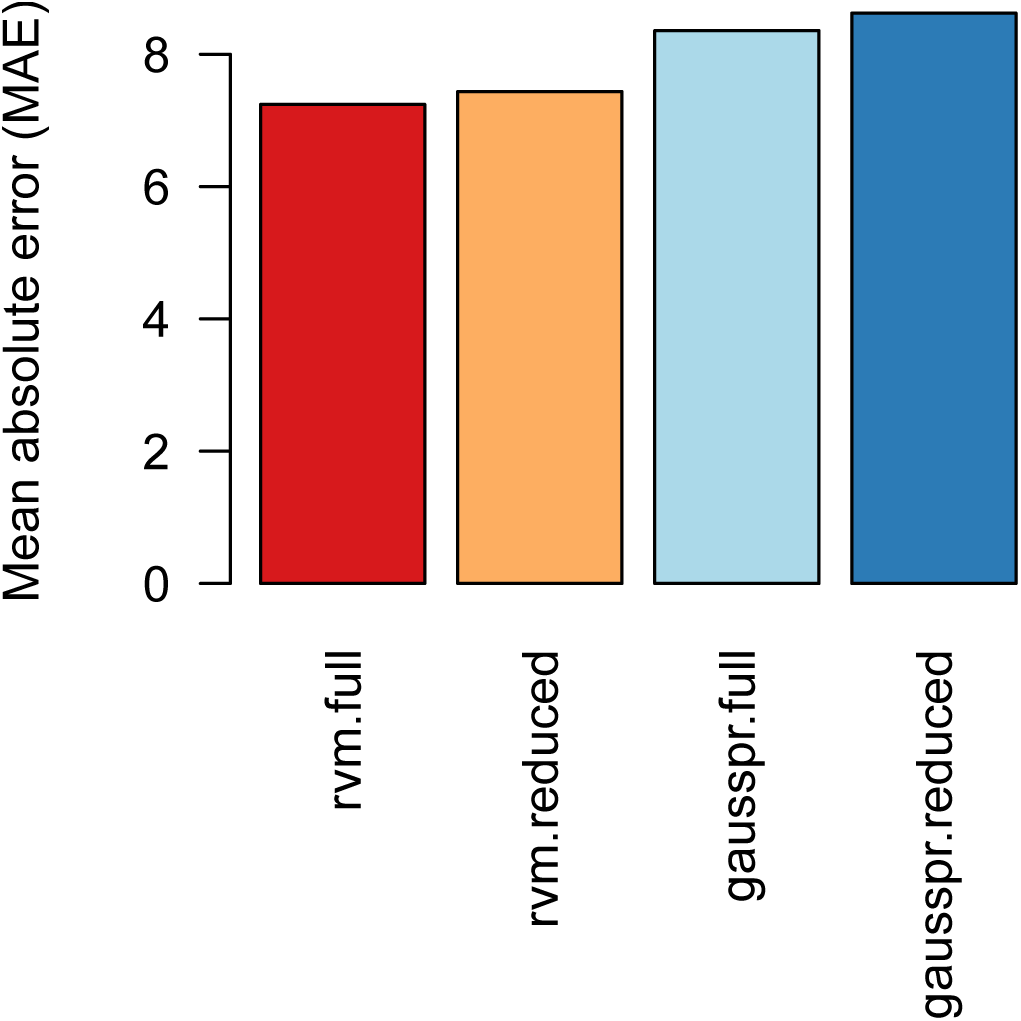
Relevance vector machine models had lower MAE compared with Gaussian Processes models. Training on the full dataset yielded small improvements in MAE for both modelling approaches.

## Discussion

This work we present NAPR, a cloud-based framework for neuroimaging-based age prediction. The approach described in this study allows external researchers to remotely access age prediction models to obtain age estimates from their own imaging data. As an example of this technique we have shared age prediction models built using machine learning techniques that are trained on cortical thickness estimates derived from Freesurfer, a commonly used and freely available morphometric analysis package. It is our hope that the NAPR approach will allow for rigorous evaluation and comparison of age prediction methods and ultimately lead to age prediction models that are accurate enough to be utilized for clinical applications. Although the NAPR system is currently designed for age prediction, it could be easily modified to provide predictions for other outcomes including the risk of developing neurological disorders.

The primary advantage of using a cloud-based framework is the ability to rapidly disseminate new predictive models. An additional benefit to providing the NAPR system as a web service is that the system is independent of the local computing setup. For example, different scanner manufacturers utilize customized compute environments. The NAPR system has minimal requirements for the end user. Essentially all that is required is a network connection, which means the system would be easy to access from any clinical imaging center. The current implementation requires imaging data to be processed using Freesurfer, however in the future image processing could be implemented on the server. Finally the general advantages associated with cloud computing apply to NAPR, including scalability, mobility and resilience against local network outages.

Results from our analyses show that our model has a higher MAE than other reported methods; our best MAE was 7.2, compared with previously reported MAEs of 1.1 - 5.8 years [1, 4, 6]. It is unclear if these differences are due to differences in modelling strategy, the input imaging data that was used to train the model, or methods for evaluating model performance. It should also be noted that there are other important metrics for evaluating the performance of age prediction models beyond mean absolute error; examples of these include model performance across age ranges, as explored in this paper, and model performance with respect to image variability (eg. motion artifact or acquisition differences). We found that there were systematic age-dependent differences in age prediction performance that were consistent across modelling techniques, which suggests that the dominant factor determining predictive model performance was our sample rather than the machine learning method used. It is likely that predictive model performance may be improved by using a standardized acquisition protocol that samples ages uniformly across the human lifespan. A potential counterargument is that training on a heterogenous multi-site dataset such as that used in our study could yield out-of-sample predictions that are fairly robust to different scanners or acquisition protocols, however external assessment is required to validate this assertion.

The overarching purpose for the development of NAPR was to provide a system for out-of-sample model testing, which is the most rigorous model evaluation technique available. By this criteria, although our model is publically available it has not yet been externally validated. Future work will address this issue. We hope that the model sharing approach advocated in this work will encourage other researchers to investigate optimal modelling approaches for estimating age using MRI data, and to share their developed models. We have provided the morphometric estimates and code that was used to train and test our model at https://github.com/hpardoe/napr. A future potential development for the NAPR platform would be to implement an automated system that allows external users to upload their own age prediction models to the server for use by the neuroscience community.

It is worth considering what would be required to improve the accuracy of MRI-based age prediction. The use of a high n, population-based collection of standardized neuroimaging data that spans most of the human lifespan would be highly beneficial for this purpose. We are not aware of the current availability of such a dataset, although it appears that large scale studies of this type are currently underway eg. the UK Biobank study [21]. Recent work has shown that multimodal imaging improves the accuracy of age prediction [1], suggesting that a comprehensive image acquisition protocol may be useful for improved age estimation methods. However since age prediction does not necessarily depend on the generation of an image, it is possible that alternative and potentially faster MR-based techniques could be developed for estimating brain age. Such a technique would be valuable for clinical applications, since patients find it difficult to tolerate longer scans. Recent work by ourselves and others have shown that participant motion during an MRI scan both (i) changes with age and (ii) systematically affects morphometric estimates [22]. Because the most clinical benefit would likely be obtained from age assessment based on neuroanatomy or brain function rather than head motion, controlling for motion via quality assurance, statistical methodology or the use of advanced image acquisitions will improve the clinical utility of neuroimaging-based age prediction methods.

An underlying assumption for the clinical utility of MRI-based age prediction is that age estimates based on structural or functional neuroimaging provides additional information above an individual’s chronological age. Furthermore many potential clinical applications of age prediction are predicated on the idea that significant differences between brain age and chronological age indicate the risk of developing a neurological disorder. Because we do not know the health outcomes for the subjects used to train the models in our study, we do not know if a high similarity between chronological and predicted age using our provided models is clinically informative. For these reasons the use of chronological age, as measured from healthy controls obtained from a diverse collection of databases, as a standard to compare with brain age may be of limited utility. Future studies could address this limitation by comparing MRI-based age estimates with alternative methods for biological age assessment (see [26] for examples of potential biological age markers), as well as tracking future health outcomes in individuals used to train the predictive model. Similarly the definition of ‘healthy’ or ‘normal’ subjects is a conceptually tricky issue that is beyond the scope of this paper.

We consider the approach described in this work as an extension of the data sharing approach that has become popular in neuroimaging research in recent years. Although there are many studies that investigate the use of neuroimaging for predictive modelling of disease outcomes or related indices in individual subjects (for recent examples see Neuroimage vol 145 Part B), traditional practice is to share summaries of the performance of these models via publication or, at best, provision of data and code. The NAPR approach provides a convenient interface for external users to access and evaluate neuroimaging-based predictive models. It is our hope that this development will encourage the widespread use and rigorous assessment of predictive models derived from neuroimaging data, and therefore assist in the translation of novel imaging methods from research laboratories to the clinic.

## Acknowledgements

The NAPR project was supported by FACES (Finding a Cure for Epilepsy and Seizures) and Amazon Web Services Cloud Credits for Research.

